# Conformational dynamics at microsecond timescale in the RNA-binding regions of dsRNA-binding domains

**DOI:** 10.1101/797449

**Authors:** H Paithankar, J Chugh

## Abstract

Double-stranded RNA-binding domains (dsRBDs) are involved in a variety of biological functions via recognition and processing of dsRNAs. Though the primary substrate of the dsRBDs are dsRNAs with A-form helical geometry; they are known to interact with structurally diverse dsRNAs. Here, we have employed two model dsRBDs – TAR-RNA binding protein and Adenosine deaminase that acts on RNA – to understand the role of intrinsic protein dynamics in RNA binding. We have performed a detailed characterization of the residue-level dynamics by NMR spectroscopy for the two dsRBDs. While the dynamics profiles at the ps-ns timescale of the two dsRBDs were found to be different, a striking similarity was observed in the μs-ms timescale dynamics for both the dsRBDs. Motions at fast μs timescale (*k_ex_* > 50000 s^−1^) were found to be present not only in the RNA-binding residues but also in some allosteric residues of the dsRBDs. We propose that this intrinsic μs timescale dynamics observed independently in two distinct dsRBDs allows them to undergo conformational rearrangement that may aid dsRBDs to target substrate dsRNA from the pool of structurally different RNAs in cellular environment.

**Statement of Significance:** This study reports for the first time the detailed characterization of microsecond timescale dynamics observed in RNA-binding regions of two distinct double-stranded RNA-binding domains (dsRBDs) using NMR relaxation dispersion experiments. dsRBDs have been known to target topologically distinct dsRNAs. However, the mechanistic details of the structural adaptation of proteins is not fully understood. We propose that the presence of such dynamics may have large-scale implications in understanding the RNA recognition mechanisms by the dsRBDs.

## Introduction

Protein-RNA interactions play a significant role in the day-to-day cellular functions. For example, the interactions between double-stranded RNA (dsRNA) and dsRNA-binding domains (dsRBDs) are involved in complex cellular processes like RNA splicing (1), RNA editing (2), RNA maturation (3), RNA transport across cellular membranes (4), etc.; and hence play a central role in regulating and executing major cellular pathways governing life. Structurally, dsRBDs contain α_1_-β_1_-β_2_-β_3_-α_2_ fold spanning a length of about 65-70 amino acid (5–7). Three regions of the dsRBD namely, 1) middle of helix α_1_, 2) loop between β_1_ and β_2_, and 3) N-terminal residues of the helix α_2_ are involved in the interactions with the backbone of the dsRNA at its minor-major-minor grove spanning a length of ~12 bp. Most dsRBDs are known to interact with dsRNAs having A-form helical structure independent of the sequence (7, 8). However, the target dsRNAs often have structural defects like internal mismatches/loops and bulges perturbing the A-form helical shape of the RNA. Such defects also lead to change in the spread of the minor-major-minor groves of the dsRNA (each represented by a unique structure defined by Euler angles (9)), thereby affecting the interaction between the dsRBD and dsRNA. Acevedo *et al.* have shown that interaction between dsRBD2 of TRBP does not lead to a significant conformational change in the substrate dsRNA (10). Thus, to effectively and efficiently target a dsRNA from the wide pool of structurally different dsRNAs in the cellular matrix, dsRBDs must adapt themselves to accommodate the conformational heterogeneity of the target dsRNAs.

The structural adaptations required in the proteins to perform a particular function involve conformational exchange processes characterized by motions at multiple timescales, including μs-ms timescale (11–14). These structural adaptations involve domain motions (15–17), catalytic interactions (18, 19), allosteric effects (20, 21), etc. Koh *et al.* have shown involvement of dynamics at the dsRNA-dsRBD interface, where the diffusion of dsRBDs of TRBP along pre-miRNA favors its cleavage to the miRNA:miRNA* duplex by Dicer – an RNase III enzyme (22). Heber et al. have recently compared the binding affinities and binding stabilities of various dsRBDs in a multi-domain protein for RNA-recognition – Staufen in *Drosophila* (23). Fareh *et al.* have shown that the recognition of target dsRNA by TRBP involves dual binding modes to discriminate between the target dsRNA and other cellular dsRNAs to which protein is exposed (24).

NMR spectroscopy can provide detailed insights into such dynamic processes at an atomic resolution (11, 25–28). Literature available till date discusses dynamics studies by NMR spectroscopy on the dsRBDs that have been restricted to the characterization of fast (ps-ns) timescale motions in these proteins and a presence of *R*_ex_ calculated using model-free analysis of such motions (29–32). Although Chiliveri *et al*. have detected slow μs-ms motions (using CPMG relaxation dispersion experiments) in the α_1_ helix of dsRBD1 of DRB4 protein of *A. thaliana* (33); they only have described *R*_*ex*_ values in their study and the detailed characterization of dynamics parameters including *p*_B_ (population of excited state), *Δω* (chemical shift difference between ground and excited state), and *k_ex_* (exchange rate between ground and excited state) values were not reported. Apart from this study, no literature thus far reports on the comprehensive characterization of μs-ms timescale dynamics in a dsRBD.

The current study attempts to characterize detailed dynamics of two model dsRBDs – namely dsRBD1 of human TAR RNA Binding Protein (TRBP2) and *Drosophila* Adenosine Deaminase Acting on RNA (dADAR) protein – at multiple timescales using NMR spectroscopy with nuclear spin relaxation and relaxation dispersion experiments. TRBP, first identified as a protein involved in an interaction with HIV TAR RNA (34), is also known to assist Dicer (an RNase III endonuclease) in processing pre-miRNAs for the generation of mature microRNAs (35–37). Being an intricate part of the canonical microRNA biogenesis pathway, TRBP encounters a wide array of structurally different dsRNAs. In addition to canonical α_1_-β_1_-β_2_-β_3_-α_2_ secondary fold, an additional N-terminal helix α_0_ was recently detected at the N terminus of dsRBD1 of TRBP2 (isoform1 of TRBP) (38, 39), the function of which remains to be understood. The second model protein used for the current investigation is the ADAR protein, known to catalyze the conversion of Adenosine to Inosine (A → I), which is critical for regulating gene expression (40–42). Human ADAR2 protein is reported to interact with dsRNAs in a sequence-specific manner (2, 43), while *Drosophila* ADAR-dsRBD1 is reported to have less sequence-specificity (41). While both the proteins contain multiple dsRBDs, this study focusses on the dsRBD1 of these proteins to allow understanding of the intrinsic dynamics features of single dsRBDs (keeping any other possible domain-domain interaction aside). We observe that the two dsRBDs have a different dynamics profile at ps-ns timescale. Interestingly, their dynamics profile at fast μs timescale is similar and is present not only in the RNA-binding regions but also in few allosteric residues of the dsRBDs. Being a common feature observed in two dsRBDs from two different proteins from two different species, we propose that these slow timescale motions are an intrinsic feature of the dsRBDs that allow them to access the ensemble of conformations, required to interact with topologically different RNAs in the cellular environment.

## Materials and Methods

### Protein Expression and Purification

The plasmids containing the gene for TRBP2-dsRBD1 and dADAR-dsRBD1 were kindly provided by Prof. Jennifer Doudna (36, 44) and Prof. Frederic Allain (41), respectively. The recombinant proteins – TRBP2-dsRBD1 and dADAR-dsRBD1 – were expressed and purified as described elsewhere (39, 41). For ^15^N labeling, the protein was prepared by growing the cells in the M9 media containing ^15^NH_4_Cl (Cambridge Isotope Laboratories) as a sole source of nitrogen.

### Circular Dichroism Spectroscopy

All the Circular dichroism (CD) experiments were carried out on Jasco J-815 CD spectropolarimeter using a quartz rectangular cuvette of 2 mm path length. The bandwidth was set at 1 nm and the data was acquired with a scanning speed of 200 nm/min with an averaging time of 1 sec. The protein samples were exchanged with Buffer A (10 mM Sodium phosphate, 100 mM NaCl, 1 mM EDTA and 1 mM DTT) pH 6.4. The CD data was acquired on 20 μM of protein in the far-UV region from 200 to 250 nm in steps of 1 nm measured in temperature range from 10 to 80 °C at steps of 5 °C. An average of 3 scans was taken at each temperature. Protein was equilibrated for at least 10 min at a given temperature before starting the measurement. All CD spectra were baseline corrected by buffer.

The CD data thus obtained were smoothened using a five-point averaging method. The fraction folded (*α*) at each temperature was calculated as (45):

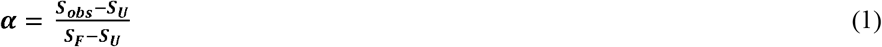

where, *S*_*obs*_, *S*_*U*_, *S*_*F*_ are the molar ellipticity at a given temperature, for the completely unfolded state and for the folded state, respectively. The thermodynamic parameters of the unfolding event were calculated by fitting CD data to Gibbs-Helmholtz equations (45):

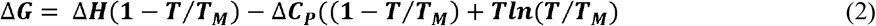

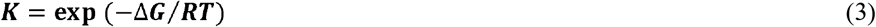

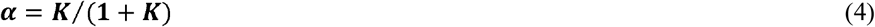

where, *ΔH* describes the enthalpy change, *ΔC_P_* the heat capacity change, *T_M_* is the temperature at mid-point of the unfolding transition, *ΔG* is the free energy of unfolding at Temperature *T* (Kelvin), *K* is the equilibrium constant of folding at temperature *T*, and *R* is the gas constant (1.98 cal K^−1^mol^−1^).

### Size Exclusion Chromatography – Multiple Angle Light Scattering

SEC-MALS analysis of the purified proteins was performed on Superdex 75 10/300 GL column (GE Healthcare) using an Agilent HPLC system equipped with the 18-angle light scattering detector (Wyatt Dawn HELIOS II) and a refractive index detector (Wyatt Optilab T-rEX). The system was calibrated with a 100 μl injection volume of 30 μM Bovine Serum Albumin solution (ThermoScientific). 100 μl protein sample was injected (in duplicates) at a concentration of 459 μM and 1.1 mM for TRBP2-dsRBD1 and dADAR-dsRBD1, respectively. The molecular weights of the peaks were determined using Zimm model in ASTRA software version 6.1.7.17 (Wyatt Technologies).

### NMR spectroscopy

All the NMR experiments were carried out on 1) Bruker 600 MHz NMR spectrometer equipped with quadruple-resonance (^1^H/^15^N/^13^C/^31^P) 5 mm Cryoprobe, X, Y, Z-gradients, and dual receiver operating; and 2) Bruker 750 MHz NMR spectrometer equipped with TXI probe, Z-gradient, and deuterium decoupling. All the NMR experiments were recorded at 25 °C. All the NMR spectra were processed using NMRPipe/NMRDraw (46) and were analyzed using SPARKY(47). Representative ^1^H-^15^N-HSQC spectra for TRBP2-dsRBD1 and dADAR-dsRBD1 with the resonance assignments (as reported in BMRB with accession number 27262 (39) and 17936 (41), respectively) have been depicted in Figure S6.

Nuclear spin relaxation experiments were measured at two field strengths (an in-house 600 MHz NMR spectrometer and 750 MHz NMR spectrometer located at IIT Bombay) on a 1.8 mM ^15^N-TRBP2-dsRBD1 sample and 0.69 mM ^15^N-dADAR-dsRBD1 sample in Shigemi tubes. For TRBP2-dsRBD1, ^15^N transverse relaxation rates (*R_2_*) were measured with eight CPMG delays of 17, 34*, 51, 68, 85, 102, 136*, and 170 ms with a CPMG frequency of 59 Hz. ^15^N longitudinal relaxation rates (*R_1_*) were measured with ten inversion recovery delays of 10, 30, 50*, 100, 200, 300, 450, 600, 750*, and 900 ms. For dADAR-dsRBD1, seven CPMG delays of 17, 34, 68*, 102, 136, 170, and 204* ms with a CPMG frequency of 59 Hz and eight inversion recovery delays of 10, 30, 50*, 100, 200, 350, 500*, and 750 ms were used to measure *R_2_* and *R_1_* relaxation rates, respectively. Delays marked with an asterisk in both the experiments were measured in duplicate for estimation of errors in relaxation rates. Steady-state [^1^H]-^15^N heteronuclear nOe measurements were carried out with a ^1^H saturation time of 3 s and a relaxation delay of 2 s for both the proteins. For the experiment without ^1^H saturation, the relaxation delay of 5 s was used. All the nuclear spin relaxation experiments were measured in an interleaved fashion and with randomized order of delays.

^15^N relaxation dispersion experiments were measured using constant time CPMG (Carr-Purcell-Meiboom-Gill) experiments (48) at two static magnetic fields (600 MHz and 750 MHz). For TRBP2-dsRBD1, each relaxation dispersion profile was composed of 9 points with *ν*_*cpmg*_ values of 25, 50*, 100, 200, 350, 500, 650*, 800, and 1000 Hz at constant relaxation time, *T_relax_*, of 40 ms. For dADAR-dsRBD1, relaxation dispersion profile was recorded with *ν*_*cpmg*_ of 50, 100, 150*, 200, 300, 450, 600*, 800 and 1000 Hz with *T*_*relax*_ of 20 ms. Duplicate points were measured for error analysis and have been marked with an asterisk.

Heteronuclear Adiabatic Relaxation Dispersion (HARD) experiments were performed at 600 MHz NMR spectrometer. A composite adiabatic pulse of 16 ms containing four hyperbolic secant family pulses of 4 ms each was used for spin-locking the ^15^N magnetization in both *R*_*1ρ*_ and *R*_*2ρ*_ experiments, as described earlier (49, 50). Relaxation dispersion was created with adiabatic hyperbolic secant pulses with different stretching factors (n=1,2,4,6,8). As the stretching factor was increased from 1 to 8, the effective spin-lock field strength increased. Relaxation delays for adiabatic *R*_*1ρ*_ and *R*_*2ρ*_ experiments were varied by varying the number of composite pulses applied during evolution. The relaxation delays used were 0, 16, 32, 48 and 64 ms corresponding to the number of composite adiabatic pulses of 0, 1, 2, 3, 4. *R*_*1*_ experiments were acquired in the same way as *R*_*1ρ*_ and *R*_*2ρ*_ experiments without using the adiabatic pulse during evolution. Relaxation delays used for the *R*_*1*_ experiment were 16, 48, 96, 192, 320, and 480 ms. In the case of *R*_1_ for HARD experiments, the delays are defined as loop of a fixed recovery delay of 16 ms (which is the time of application of adiabatic pulses in *R*_1*ρ*_, and *R*_2*ρ*_ experiments). An inter-scan delay of 2.5 s and 3 s was used for TRBP2-dsRBD1 and dADAR-dsRBD1, respectively in all the above relaxation experiments.

### NMR relaxation Data Analysis

Relaxation rates in *R*_*1*_/*R*_*2*_/*R*_*1ρ*_/*R*_*2ρ*_ experiments were calculated by fitting the intensity data against relaxation delays to mono-exponential decays in Mathematica (51). Errors in the relaxation rates were calculated as fit errors using a combination of duplicate delay and Monte Carlo simulations. [^1^H]-^15^N nOe values were obtained as a ratio of the intensity of respective peaks of the spectra recorded with and without saturation. Errors in nOe values were obtained by propagating the errors from RMSD values of baseline noise as obtained from Sparky.

Analysis of the ^15^N-relaxation data (*R*_*1*_, *R*_*2*_, [^1^H]-^15^N nOe) recorded at two magnetic fields was done using the extended model-free formalism (52–55) with the graphical user interface provided by the Relax v4.0.3 software (56, 57). The ^1^H-^15^N dipole-dipole interaction and CSA values (−172 ppm) were included in calculations as per the default protocol. The distance between N and H was set as 1.02 Å. First, the local *τ*_*m*_ diffusion model was optimized with no global diffusion parameter defined. Diffusion tensor values for various diffusion models – sphere, spheroid, and ellipsoid – were calculated using the program quadric diffusion (58) for data recorded at both the magnetic fields. Solution structure of TRBP2-dsRBD1 (39) and dADAR-dsRBD1 (41) was used to optimize the diffusion tensor parameters. These diffusion tensor parameters were then used to calculate the model-free parameters from the ten model-free models and the best model for each residue was selected based on Akaike’s Information Criteria. After the selection of these local models, diffusion tensor was further optimized with fixed local models. This was repeated until all the model-free parameters converged. The global diffusion model was then selected based on the chi-squared values obtained from optimized diffusion models. The error analysis for the selected model (global and local) was done using the Monte Carlo method.

For CPMG relaxation dispersion experiments, effective transverse relaxation rates at each CPMG frequency were extracted using the following equation (27):

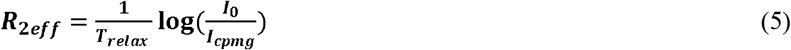

where, *I*_*0*_ is the intensity of a peak in reference spectrum and *I*_*cpmg*_ is the intensity of the respective peak in the spectrum with CPMG field applied at a frequency *ν*_*cpmg*_. The *R*_*2eff*_ rates thus obtained were plotted against the CPMG frequency *ν*_*cpmg*_.

For HARD experiments, the relaxation rates (*R*_*1*_, *R*_*1ρ*_, and *R*_*2ρ*_) along with rotational correlation time calculated from the model-free analysis was used to fit the data to the time-dependent Bloch-McConnell equation. Since the general solutions to the equations describing the evolution of bulk magnetization in chemical exchange during adiabatic relaxation process are not available, a numerical solution was used by using a geometric approximation method to extract the exchange parameters from the Bloch-McConnell equation (59). The parameters that describe these motions were extracted under the assumption of a two-site exchange model. The relaxation rates (*R*_*1ρ*_ and *R*_*2ρ*_) have been obtained by using a train of hyperbolic secant family adiabatic pulses during evolution of the magnetization. Further, the shape of these pulses are changed by changing the stretching factor of the pulse to create dispersion in the *R*_*1ρ*_ and *R*_*2ρ*_ rates. Libraries of solution points of the Bloch-McConnell Equation were kindly provided by Dr. Fa-An Chao, which were created by assuming that the *R*_*1ρ*_ and *R*_*2ρ*_ rates are the linear combinations of the functions of *R*_1_, *R*_2_, and *R*_ex_ for the defined set of dynamic parameters (*k*_ex_, p_A_, Δω, and offset). A rigorous grid search through these solution surfaces by Monte Carlo method helped to extract the exchange parameters that best fit the observed dispersion in *R*_*1ρ*_ and *R*_*2ρ*_ rates obtained from use of various shapes of the adiabatic pulses. The solution to the equation by using a geometric approximation method to extract the dynamic parameters and the relaxation rates during the chemical exchange process has been described in detail elsewhere (59).

## Results

### Thermal melting profiles of the two dsRBDs

To investigate the thermal stability of the dsRBDs, far-UV CD-based melting studies were carried out. The temperature-dependent far-UV CD spectra for TRBP2-dsRBD1 showed that the protein conformation starts to open at 25°C (where all the measurements are carried out in this study) (Figure 1). The unfolding event was characterized by an enthalpy change (*ΔH*) of 41.11±2.32 kcal/mol and corresponding *T*_*M*_ of ~44°C obtained by fitting of the CD data with Gibbs-Helmholtz equation (see Methods section). The temperature-induced unfolding of dADAR-dsRBD1 using far-UV CD showed a very similar profile with enthalpy change (*ΔH*) of 33.58±1.57 kcal/mol and corresponding *T*_*M*_ of ~44°C (Figure 1). Though the thermodynamics parameters, obtained from Gibbs-Helmholtz equation, cannot be correctly interpreted due to irreversible nature of the unfolding transition (45), the melting profiles with low *T*_*M*_ values (similar values were obtained from the derivative plot dθ/dT vs Temperature, Figure 1E,F) hinted towards low thermal stability of the proteins at 25°C.

**Figure 1:**
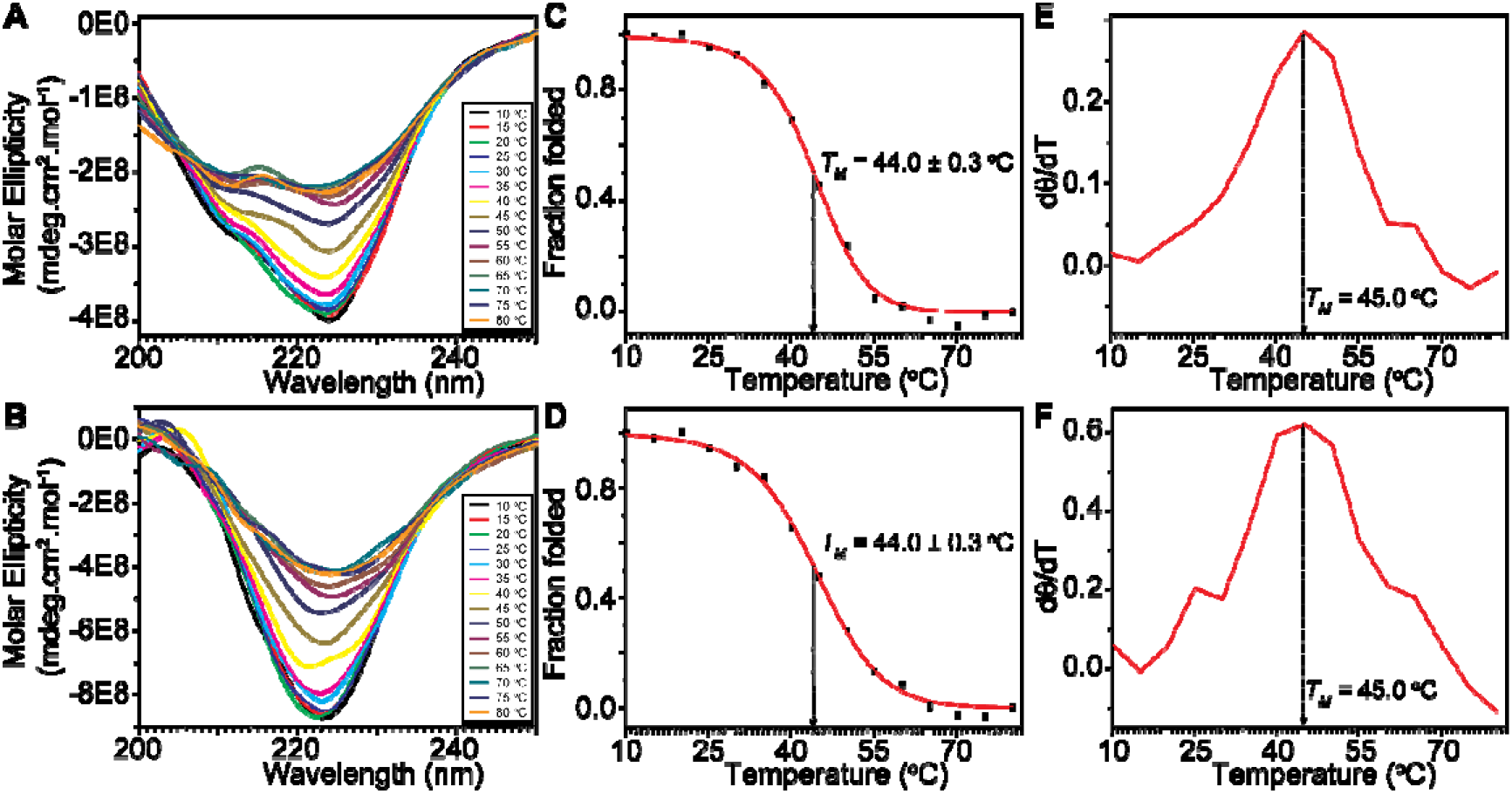
Thermal melting profiles of the two dsRBDs by Circular Dichroism. The normalized far-UV CD spectra of (**A**) TRBP2-dsRBD1 and (**B**) dADAR-dsRBD1 where molar ellipticity is plotted against wavelength measured as a function of temperature from 10 to 80 °C at an interval of 5 °C. Inset shows a color scheme for the data measured at various temperatures. Fraction folded of (**C**) TRBP2-dsRBD1 and (**D**) dADAR-dsRBD1 calculated from ellipticity at 222 nm using equation 1 plotted against temperature to highlight the melting profile of the protein. First derivative plot (dθ/dT vs Temperature) of the ellipticity at 222 ns for (**E**) TRBP2-dsRBD1 and (**F**) dADAR-dsRBD1. The maxima of the curve represent the melting temperature (*T*_*M*_).

### TRBP2-dsRBD1 and dADAR-dsRBD1 show differential dynamics at the ps-ns timescale

The spectral density function, J(ω), used to describe motions by virtue of time correlation function of the bond vector can be analysed by a variety of methods, including, phenomenological interpretation of the relaxation variables, spectral density mapping, model-free analysis, and model-dependent analysis (11, 60). The phenomenological interpretation of the site-specific relaxation variables (*R*_1_, *R*_2_, and [^1^H]-^15^N nOe) remains limited towards assigning the structural details to the motions. The model-dependent analysis used to be a method of choice before the introduction of the Lipai-Szabo model-free approach (52, 53). Although specific models are highly useful, but typically due to unavailability of detailed side-chain data for proteins, model-free and spectral density mapping became the methods of choice after they are introduced. The spectral density approach, pioneered by Peng and Wagner (61), gives a great deal of information about the site-specific spectral density functions defining the NMR relaxation phenomenon. The spectral density mapping approach, however, does not consider the global motions of the protein that might be present in the structured proteins. Spectral density mapping also does not contain any parameter to define the motional amplitude. Model-free approach defines the spatial restriction of the bond-vector by a generalized order parameter approach for isotropic and anisotropic systems in addition to global and local effective correlation times. While model-free approach also uses spectral density functions to fit the relaxation data, it leaves S^2^ and τ_e_ as adjustable parameters while data fitting. To investigate the timescale of motions present in the proteins, NMR spin relaxation data (*R*_*1*_, *R*_*2*_ and [^1^H]-^15^N NOE) was analysed for 69 non-overlapping peaks out of 96 assigned peaks for TRBP2-dsRBD1 (39) and for 58 such peaks out of 74 assigned peaks for dADAR-dsRBD1 (41) (Figure S1). The analysis of the data using the extended model-free approach (52–55) allowed the extraction of dynamic parameters at the ps-ns timescale.

The order parameter *S*^*2*^ plotted against the residue number for TRBP2-dsRBD1 have been depicted in Figure 2A,B. The order parameters (S^2^) calculated from the model-free analysis are interpreted most-frequently as diffusion in a cone model. In this model (53, 54), the N-H bond vector is assumed to freely diffuse within a cone defined by the semi-angle θ such that:

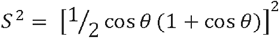

**Figure 2:**
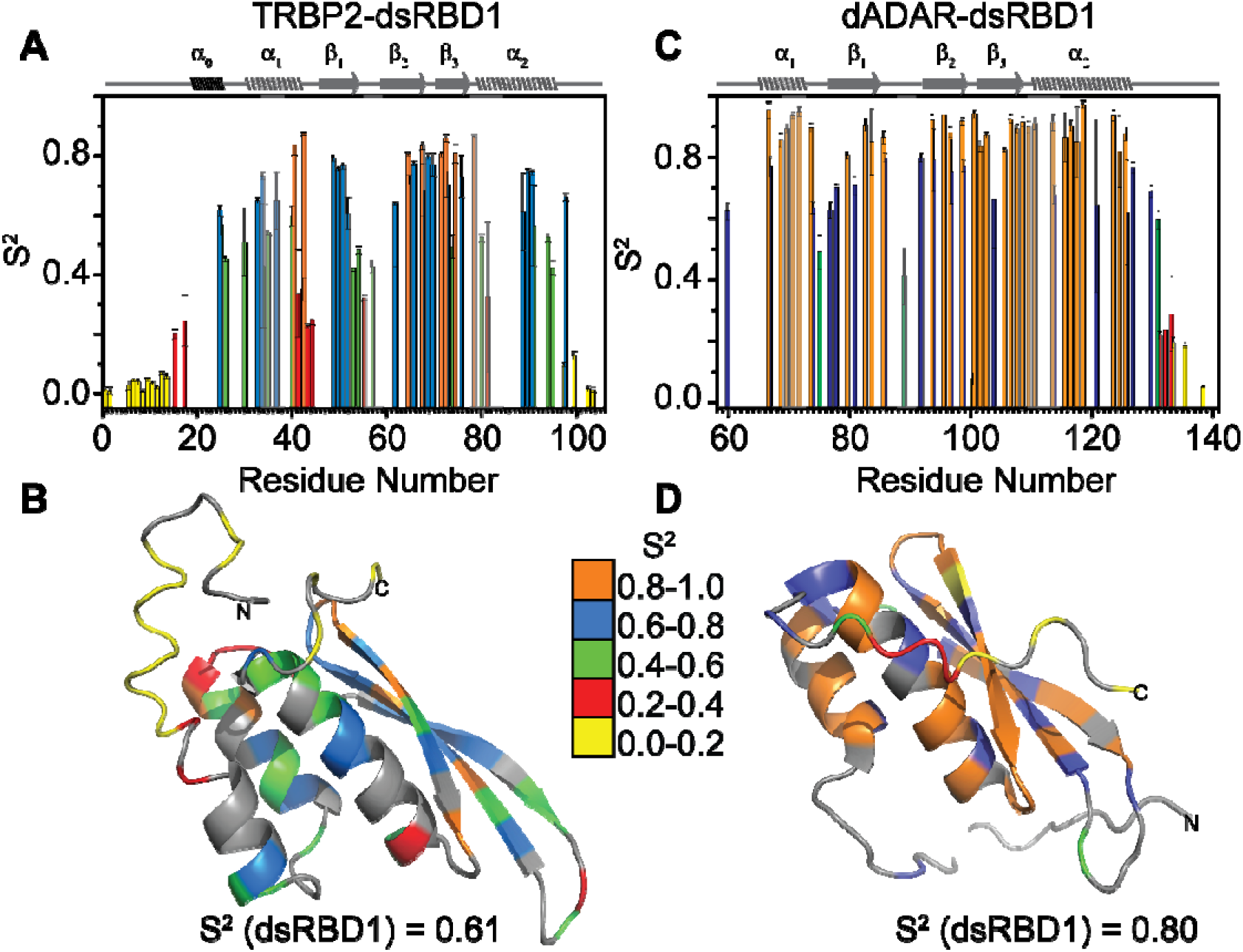
Differential dynamics at ps-ns timescale in the two dsRBDs. Order parameters (S^2^) calculated from the extended model-free approach plotted against residue number for (A) TRBP2-dsRBD1 and (C) dADAR-dsRBD1; *S*^*2*^ values mapped in color codes on the tertiary structure of (B) TRBP2-dsRBD1 and (D) dADAR-dsRBD1. The secondary structure of the protein has been shown on the top of the section (A) and (C); and color codes for the *S*^*2*^ values have been mentioned alongside section (D). Average *S*^*2*^ values (excluding the N- and C- terminal flexible regions) for both the proteins have been depicted at the bottom.

The smaller cone angle represent restricted motions with S^2^ value close to 1 and mobile residues that can swap larger cone angle have S^2^ value close to 0. Thus, the residues with lower S^2^ value in the stretches near RNA-binding residues represent motion in cone with wide semi-angle and thus reflect higher amplitude of motion. The plot showed large variations of *S*^*2*^ values throughout the protein backbone. The mean *S*^*2*^ for the core RNA-binding region (31-95 aa) was found to be 0.61; which suggests the presence of fast motions at ps-ns timescale in the structured regions of the protein. The residues in the α_0_ helix (19-25 aa) of the protein showed higher *S*^*2*^ than the N-terminal unstructured region (1-16 aa) and lower *S*^*2*^ than the dsRBD core indicating the structured yet dynamic stretch of the residues in this helix. Further, the *S*^*2*^ values in the stretches near the RNA-binding region showed a higher amplitude of motions (as described above) compared to the overall motions of the dsRBD core. Around 78% of the analyzed residues of TRBP2-dsRBD1 showed the best fit to a ‘model-free’ model with motions in two timescales. The fast internal motions of the timescale of tens of picoseconds reflected random thermal motions, which were observed at the terminal and in the loop regions in the protein. Slower motions at nanosecond timescale (1-4 ns), but faster than the global tumbling time of the protein (7.64 ns), were present throughout the protein backbone.

The residues best fit with a model containing exchange term (*R*_*ex*_) indicated the presence of motions at μs-ms timescale, which involved conformational exchange processes (Figure 3A). The residues having large *R*_*ex*_ were mainly observed in helix α_1_ and α_2_ near the RNA-binding region, and the loop regions of the protein which correlated with the lower *S*^*2*^. Interestingly, a pair of residues in helix α_2_ (A88 and E89) showed high *S*^*2*^ values and *R*_*ex*_ at the same time.

**Figure 3:**
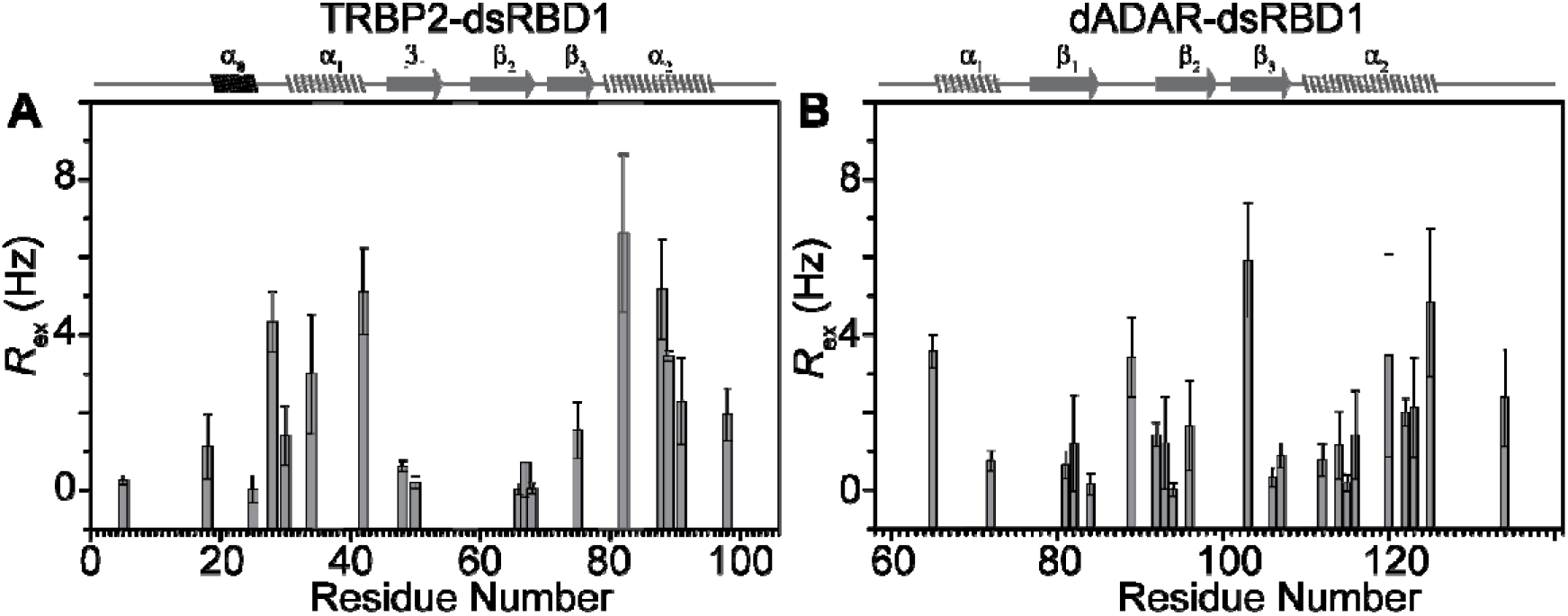
Slow μs-ms timescale dynamics pointed out by model-free analysis. *R*_*ex*_ values derived from the extended model-free approach of the ^15^N spin relaxation data (*R*_*1*_, *R*_*2*_ and nOe) for (A) TRBP2-dsRBD1 and (B) dADAR-dsRBD1 with secondary structure mentioned on top of plot and grey columns indicate RNA binding regions.

The *S*^*2*^ values calculated from model-free analysis of nuclear spin relaxation data for dADAR-dsRBD1 showed higher rigidity (when compared with that of TRBP2-dsRBD1) in the structured region of the protein with an average *S*^*2*^ of 0.80 (Figure 2C,D). The only residues that showed more flexibility with lower than average *S*^*2*^ value were the loop regions. The residues in the protein backbone showed motions at ns timescale (~2 ns) with fast motions at ps timescale in loop residues. Interestingly, many residues near RNA-binding region (T65, L72, E81, S82, T84, F92, I94, Q106, G107, E116, L122, R123, and S124) showed the presence of μs-ms timescale motions (similar to what was observed in TRBP2-dsRBD1) as comprehended from *R*_*ex*_ terms (Figure 3B) calculated by the extended model-free analysis. The global tumbling time for the dADAR-dsRBD1 was estimated to be a 9.39 ns from the model-free analysis.

### Conformational exchange in the μs-ms timescale in dsRBDs

Driven by the presence of *R*_ex_ along the length of the two dsRBDs, dynamics at μs-ms timescale was tested using relaxation dispersion experiments. It is important to note here that although the *R*_ex_ values derived from the spin-relaxation data and model-free analysis suggested a presence of slow motions in the two dsRBDs. This, however, may fail to provide a complete picture of the microsecond timescale dynamics mainly due to the fact that in such an experimental setup, *R*_2_ is measured only on a single Carr-Purcell-Meiboom-Gill (CPMG) frequency on two different magnetic fields. A complete picture is obtained recording by the relaxation dispersion experiments (CPMG or *R*_1*ρ*_) where relaxation rates are measured as a function of CPMG frequencies or the spin-lock field strength. The dynamics measurements at μs-ms timescale were initially carried out by using CPMG relaxation dispersion experiments (11, 27). However, *R*_*2eff*_ was found to be invariant to the CPMG frequencies (Figure S2 and Figure S3) used to create the dispersion thereby suggesting that the dsRBDs are not conformationally exchanging at the timescale (~0.3 to 10 ms (11)) sensitive to CPMG-RD experiments.

Next, dynamics at the μs-ms timescale was tested using Heteronuclear Adiabatic Relaxation Dispersion (HARD) (49, 50, 59) NMR experiment that monitors NMR spin relaxation in a rotating frame in the presence of hyperbolic secant (HSn, where n = stretching factor) adiabatic pulses. HARD NMR experiment allows to explore the conformational exchange processes occurring on the 10 μs to 10 ms timescales. The plot of relaxation rates *R*_*1ρ*_ and *R*_*2ρ*_ against residue numbers showed that as the applied field strength increases from HS1 to HS8, the *R*_*1ρ*_ rates increased and the *R*_*2ρ*_ rates decreased for both the dsRBDs (Figure 4). The dispersion in the relaxation rates observed points to the dynamic processes that can be probed by this method. Assuming two-state exchange between a ground state and an excited state, fitting of the *R*_*1*_, *R*_*1ρ*_, and *R*_*2ρ*_ rates to the solution of the Bloch-McConnel equations by using geometric approximation approach (59) allowed to extract the residue-specific dynamic parameters (the conformational exchange rate, *k*_*ex*_, the equilibrium populations of the two states, *p*_*A*_ or *p*_*B*_, and the chemical shift difference, *Δω*). The two-state model is the simplest model to describe the chemical exchange in residues in the protein where the exchange is assumed to have conformations described by state A (ground state) and state B (excited state) such that the transition between state A and B occurs with exchange rates defined by *k*_ex_. Though the model describes exchange between these two states, the residues can have multiple conformations other than state A and B. Since the data could be explained by a simple model (with less number of parameters), more complex models (with higher number of parameters due to multiple states) were not tried for fitting. A more complex model (than a simple two-state model) would always fit better as it will have more number of parameters.

**Figure 4:**
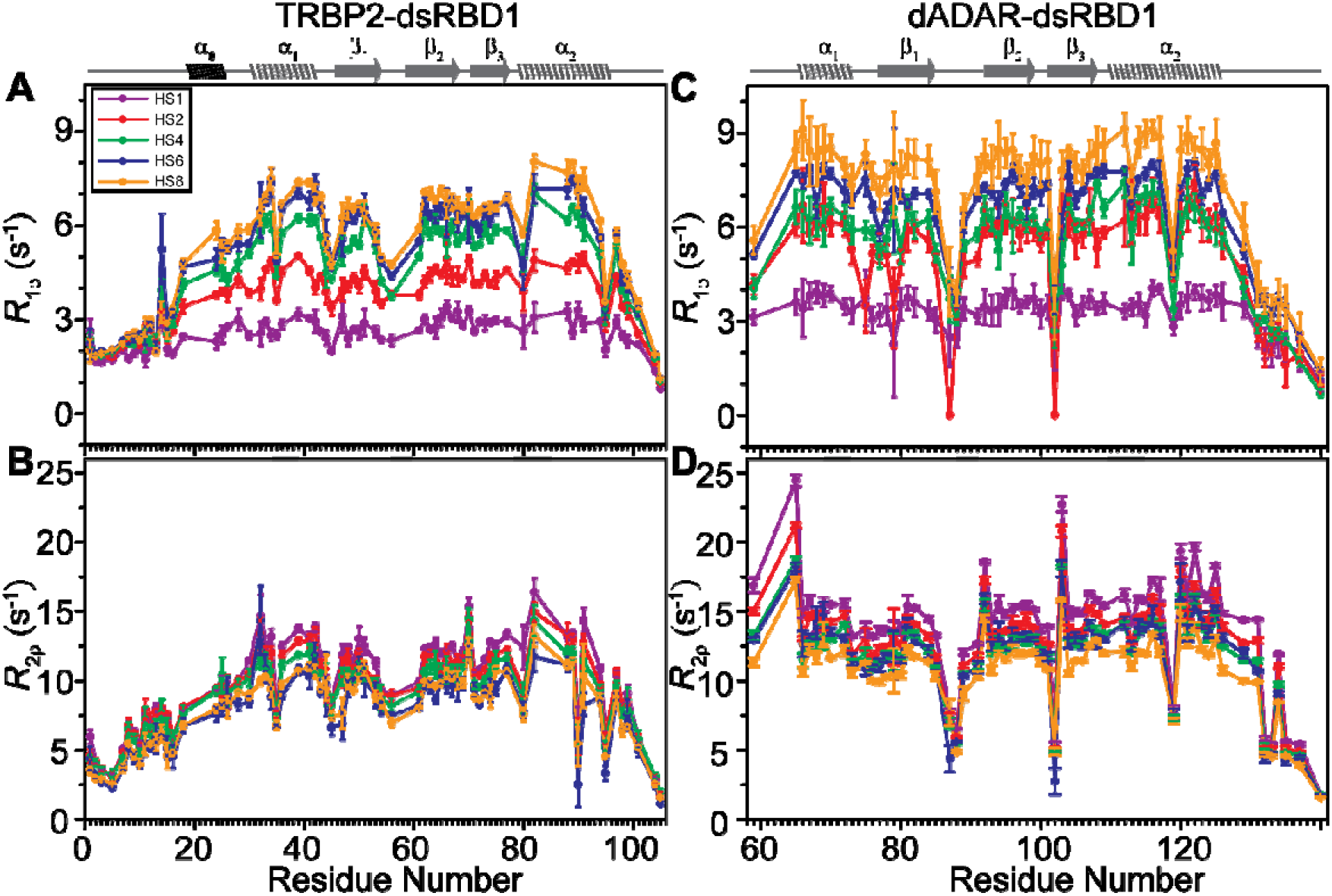
Dispersion in relaxation rates for two dsRBDs. (A) *R*_*1ρ*_ and (B) *R*_*2ρ*_ relaxation rates derived from the adiabatic relaxation dispersion experiments plotted against residue number as a function of adiabatic pulse stretching factors (color coded for n=1,2,4,6, and 8) for TRBP2-dsRBD1. (C) *R*_*1ρ*_ and (D) *R*_*2ρ*_ relaxation rates from the adiabatic relaxation dispersion experiments plotted for dADAR-dsRBD1.

Since there is a partial overlap in the timescale of motions that can be probed by CPMG-RD and HARD NMR experiments, we simulated the *R*_*2eff*_ rates for CPMG relaxation dispersion using the dynamics parameters obtained from HARD NMR experiments to verify the absence of dispersion in *R*_*2eff*_ rates as observed in CPMG-RD. The simulations for *R*_*2eff*_ values for CPMG relaxation dispersion experiment were performed using the following equation (11):

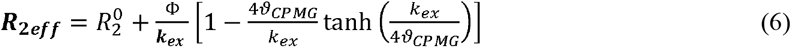

where, Φ = *p_A_* * *p_B_* * Δ*ω*^2^ with *p*_*A*_, *p*_*B*_, Δω, and *k*_*ex*_ values used in simulation were obtained from HARD NMR data analysis. υ_CPMG_ is varied in the simulations as per the range used to record CPMG-RD data.

The simulation of the *R*_*2eff*_ rates for CPMG relaxation dispersion with equation 6 with the dynamics parameters extracted from HARD data showed no dispersion in the *R*_*2eff*_ relaxation rates.

Careful analysis of the *k*_*ex*_ and the *p*_*B*_ values for TRBP2-dsRBD1, depicted on the protein structure (Figure 5A), showed that the residues in the core RNA-binding region exhibited large *k*_*ex*_ (>50000 s^−1^) with a large excited-state population (Figure S4). The L35 and Q36 residues (lie in the middle of the α_1_), the residue E54 (in the β_1_-β_2_ loop), and the residue A82 (lies towards the N-terminal of the helix α_2_) displayed large *k*_ex_ values. A few of the non RNA-binding residues (A25, N26, N61, F64, T67, and V68 – although not directly interacting with dsRNA, but are present in purlieu of RNA-binding residues) also showed *k*_*ex*_ > 50000 s^−1^. The observed exchange in these residues might have originated to maintain the overall tertiary fold of the protein during the conformational exchange happening in the RNA-binding residues. The residues in the middle of β strands, which are involved in maintaining the tertiary structure of the protein through a network of interactions (hydrophobic interactions or charge-charge interactions or H-bonding interaction) and the residues in loop/terminal regions displayed a conformational exchange occurring at the rate of 50000 s^−1^> *k*_*ex*_ >5000 s^−1^. These residues included (M1, T10, G13, C14, L51, V66, C73, H94, L95, A104) and about 10-40% of the population of these residues existed in excited states. Rest of the residues showed populations of <10% and are exchanging at rates <5000 s^−1^. Along with the wide range of *k*_*ex*_ and the *p*_*B*_ values, wide dispersion (5-1100 Hz) in the corresponding *Δω* values (between ground state and excited state) in ^15^N indicated the availability of a large conformational space which the protein may populate allowing target dsRNAs to select a conformation for efficient binding (Figure S4).

**Figure 5:**
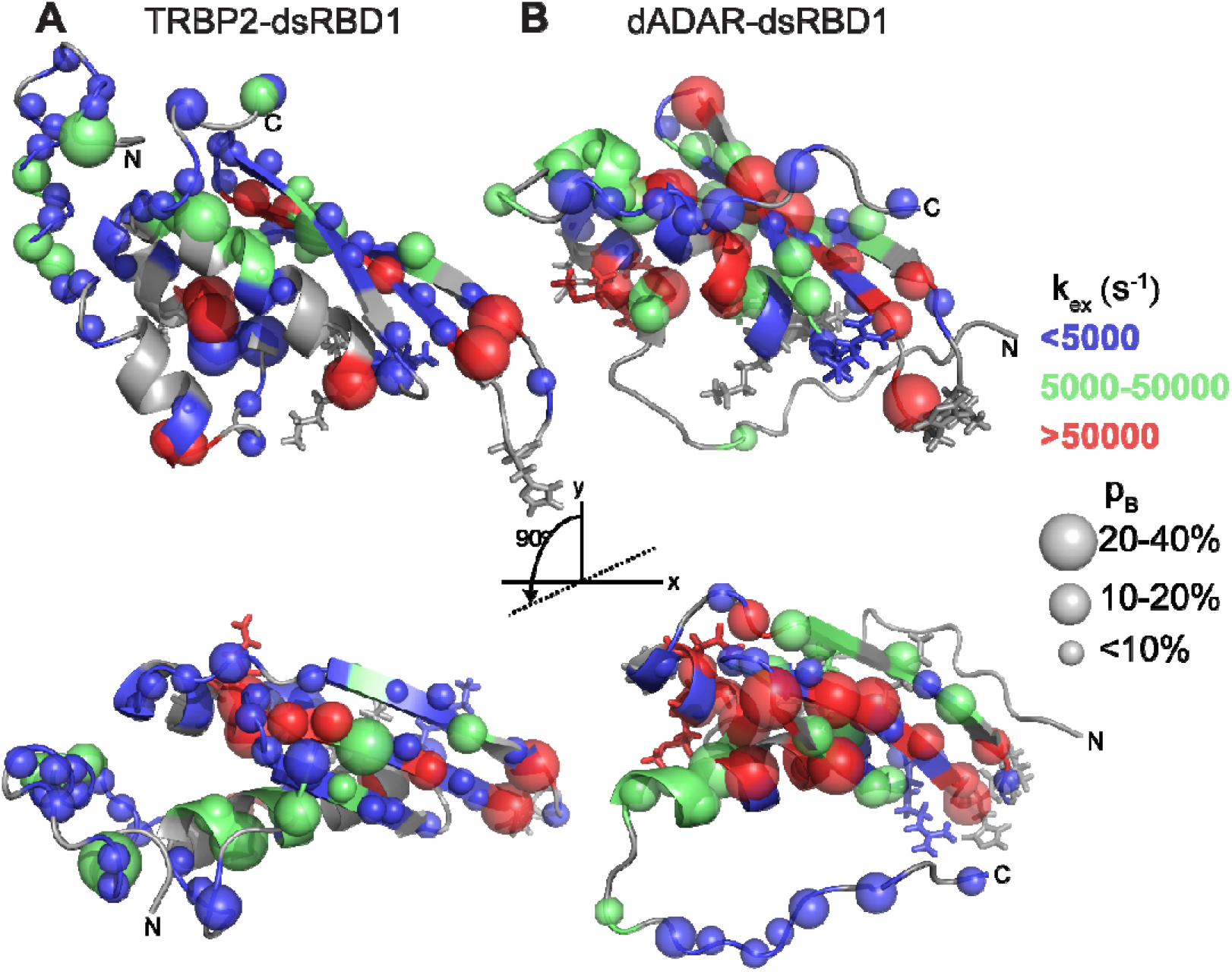
Conformational exchange along the backbone of the two dsRBDs. Mapping of dynamics parameters, rate of exchange between ground state and excited state (*k*_*ex*_), and population of the excited state (*p*_*B*_) as obtained by the ‘geometric approximation method’ from HARD experiment, on the tertiary structure of (A) TRBP2-dsRBD1 and (B) dADAR-dsRBD1. The color code has been used to highlight the distribution of *k*_*ex*_ values whereas the size of the sphere highlights the variation in the *p*_*B*_ values across protein backbone. The RNA binding residues have been shown in stick mode. The structure of the protein is shown in two orientations by 90° rotations about the x-axis in order to observe clustering of the residues with similar exchange rates at the rear face of the protein.

Despite differences in the primary sequence of the two dsRBDs, a strikingly similar behaviour of *k*_*ex*_ was observed in dADAR-dsRBD1 (Figure 5 and Figure S4). Residues lying in the core RNA-binding region of the dsRBD, including residues in the middle of α_1_ (V66, A67, M68, N70), residues in β_1_-β_2_ loop (T84, A89, L91), and residues towards N-terminal of α_2_ (R114, I115, E116, A117) were all found to have *k*_*ex*_ > 50000 s^−1^. Similar to TRBP2-dsRBD1, a few of the non-RNA binding residues (L76, T93, I94, S95, V96, Q101, and L104) also showed *k*_*ex*_ > 50000 s^−1^ in the region suggesting allosteric role (L76, I94, and L104 directly interact with RNA-binding residues) for the proposed conformational adaptation by dsRBD. Other residues in the non-RNA-binding region had *k*_*ex*_ between 5000 s^−1^ to 50000 s^−1^ and were found to lie in either loops or in the middle of β strands and the fraying end of helix α_2_. Rest of the residues showed populations of <10% and are exchanging at rates <5000 s^−1^. A similar chemical shift dispersion was observed in the *Δω* values in ^15^N for dADAR-dsRBD1 (Figure S4) as observed for TRBP2-dsRBD1.

## Discussion

In order to understand the role of protein dynamics in interaction with the topologically broad range of dsRNAs, two dsRBDs – TRBP2-dsRBD1 and dADAR-dsRBD1 – were tested for the presence of dynamics. The sequence identity between the two dsRBDs was found to be 23.6% and the similarity of 33.3% with the canonical tertiary fold common to all dsRBDs. Despite the large differences in the primary sequence, both the proteins showed a similar thermal stability as evidenced by the similar CD melting profile with a *T*_*M*_ of ~44°C (Figure 1). Fraction-folded, as calculated from the CD data for TRBP2-dsRBD1 and dADAR-dsRBD1, showed that ~95% of the population of the dsRBDs exist in folded state at 25°C, where all NMR experiments are recorded.

Interestingly, the analysis of nuclear spin relaxation experiment by extended model-free approach showed lower order parameter values (***S*^*2*^**<0.6) in the structured region of the protein in addition to the low values in the loop and terminal areas for TRBP2-dsRBD1. This supported the internal flexibility in an otherwise globular protein. Order parameters for dADAR-dsRBD1, however, showed routinely observed profile with lower values (***S*^*2*^**<0.4) in the loop and terminal regions and higher values (***S*^*2*^**>0.8) in the structured regions of the protein. The global tumbling time for the two proteins as calculated from the model-free analysis was higher (7.64 ns for TRBP2-dsRBD1 and 9.39 ns for dADAR-dsRBD1) than that observed routinely for a ~11-12 kDa protein. A higher global rotational correlation time is attributed to the presence of conformational heterogeneity, the presence of an oligomeric species, and the presence of large amount of unstructured regions in the two proteins. The presence of an oligomeric species affecting the correlation time is ruled out by size-exclusion chromatography coupled with multiple angle light scattering study of the proteins. While dADAR-dsRBD1 (rotational correlation time of 9.39 s) showed a pure monomeric species, TRBP2-dsRBD1 (rotational correlation time of 7.64 ns) showed only a minor population of tetramer (Figure S5). With a ~45 kDa molecular weight, transverse relaxation rate, *R*_2_, values of tetramer would significantly broaden the NMR peaks for tetramer, and additionally with a small population (one-fourth of that of monomer), tetramer peaks would be beyond detection in an HSQC spectrum. Also, since the two species (monomer and tetramer) co-exist in solution for TRBP2-dsRBD1, the monomer-tetramer exchange would be in slow timescale regime and thus would not affect the relaxation rates measured for monomer.

Secondary structure analysis of the TRBP2-dsRBD1 shows that it has 45% residues in the unstructured and 55% residues in the structured regions. A similar analysis showed 59% residues in unstructured and 41% residues in structured regions of dADAR-dsRBD1. Thus, the presence of unstructured regions in the two dsRBDs could be one of the reasons for relatively higher correlation times than the globular proteins of this size. A similar value of global tumbling time has been previous reported for other dsRBDs, e.g., DGCR8-dsRBD1 (7.2 ns), Drosha-dsRBD (6.29 ns), and Dicer-dsRBD (6.35 ns) (30, 31).

The presence of slow motions at μs-ms timescale suggested by model-free analysis is further tested using heteronuclear adiabatic relaxation dispersion (HARD) NMR experiment. Analysis of HARD NMR data showed large dispersion profiles in both *R*_*1ρ*_ and *R*_*2ρ*_ rates when measured as a function of adiabatic pulse stretching factor, suggesting the presence of slow-dynamics at μs-ms timescale (Figure 4). The presence of strikingly similar dispersion profile in relaxation rates in both the dsRBDs indicated parallel dynamics at μs-ms timescale. For example, residues in the core RNA-binding regions of the two dsRBDs were found to have *k*_*ex*_ > 50000 s^−1^ to excited state(s) and are characterized by a large change in ^15^N chemical shift values (*Δω*) and significant populations in excited state(s) (Figure 5 and Figure S4). This fast microsecond timescale dynamics (*k*_ex_ > 50000 s^−1^) are present in the RNA-binding residues or in their close proximity (in the middle of α_1_-helix and at the C-terminus of β_1_-strand and N-terminus of β_2_-strand), and in few other non-interacting residues located in β-strands in both the dsRBDs. The residues in the C-terminus of the α_2_-helix of the two dsRBDs, which is in proximity to the RNA-binding residues in α_1_-helix, show motions with *k*_ex_ between 5000 s^−1^ and 50000 s^−1^. Significant excited state population with a dispersion of values of *Δω* from 5-1100 Hz is suggestive of large conformational dynamics present in the two dsRBDs. In addition to the faster μs timescale motions in the RNA-binding regions, conformational exchange with slower exchange rates (*k*_ex_ < 5000 s^−1^) was found in a large number of residues (49 in TRBP2-dsRBD1 and 25 in dADAR-dsRBD1, respectively) in the non-RNA-binding regions of the proteins. Interestingly, despite the significant differences in the primary sequence of the two dsRBDs, the motions at μs-ms timescale are highly similar in RNA-binding residues and in a few allosteric regions. This suggests that the analogous dynamics observed by relaxation dispersion method in two dsRBDs are not affected by primary sequence and are intrinsic to the protein’s secondary/tertiary structure. Additionally, higher *k*_*ex*_ values observed in non-RNA-binding regions manifests in the allosteric role being played by these residues to maintain the structural integrity in the conformational exchange process (Figure 5 and Figure S4).

Since the exchange is present all along the protein backbone and the sign of *Δω* cannot be determined by the HARD experiment(59) we could not calculate the structure of the excited state(s). Further, the co-existing tetramer species in TRBP-dsRBD1 is detected (in SEC-MALS) at a much slower timescale than NMR (eluted from the column over an hour), thus, we rule out the possibility of microsecond dynamics due to the multimer formation. While no such heterogeneity is found in dADAR-dsRBD1 sample, the presence of microsecond timescale dynamics was still observed suggesting that the origin of this dynamics is due to a common process present in both the proteins. A simple unfolding of the protein with a small population (of unfolded species) at 25°C may allow it to access the energy barrier required to populate alternative conformational states and thus may lead to the observed exchange.

Here, for the first time, we report the detailed characterization of the conformational dynamics of the dsRBDs, which suggests that the presence of similar μs-ms timescale motions in two dsRBDs. Further, it hints towards these motions as a general feature of dsRBDs. These conformational dynamics lead to the creation of a conformational pool for the dsRBDs that might allow it to interact with target dsRNAs in the cellular environment. This may further allow processing of the accurate substrate dsRNAs by the catalytic protein partners of dsRBDs. Identification and characterization of the dynamics at the μs-ms timescale shows conformational adaptability of the dsRBDs and is a step towards in understanding the mechanism of non-specific interaction between dsRBD and target dsRNAs.

## Conclusions

The observations show the presence of conformational plasticity in two dsRBDs irrespective of differences in their primary sequences. The effect of different primary sequences was reflected in the ps-ns timescale dynamics of the two proteins. However, the similar *R*_*ex*_ rates measured by the extended model-free approach and the μs-ms timescale dynamics measured by the NMR relaxation dispersion method are present all along the backbone of the protein in addition to the faster μs timescale motions in the RNA-binding region. This conformational plasticity and the μs-ms timescale dynamics in two dsRBDs might be responsible for the adaptation required by the dsRBDs to target conformationally distinct dsRNAs present in the cellular pool.

## Author Contributions

J.C. and H.P. conceived the approaches to dynamically characterize the model dsRBDs and wrote the paper. H.P. performed all the experiments and the data analysis in active discussions with J.C.

## Acknowledgements

Authors thank Prof. Jennifer Doudna (University of California, Berkeley) for the TRBP plasmids and Prof. Frederic H.T. Allain (ETH Zurich) for ADAR-plasmid. Authors also acknowledge High Field NMR facility at IISER-Pune (co-funded by DST-FIST and IISER Pune) and the High-Field NMR facility at IIT Bombay. This work was supported by funding from Indian Institute of Science, Education and Research, Pune; Department of Biotechnology, Govt. of India [No. BT/PR24185/BRB/10/1605/2017]; and extramural funding from the Science and Engineering Research Board (SERB), Govt. of India [EMR/2015/001966]. We acknowledge Dr. Radha Chauhan, Ms. Sangeeta Niranjan, and Ms. Bhawana Burdak for the help with measurement and analysis of the SEC-MALS data at NCCS, Pune. Authors also acknowledge Prof. Gianluigi Veglia (University of Minnesota, Minnesota) and Dr. Fa-An Chao (National Cancer Institute, Maryland) for active discussions while setting up HARD experiments and data analysis. Authors thank Dr. Arnab Mukherjee and Dr. T.S. Mahesh at IISER Pune for helpful discussion as Research Advisory Committee members for HP. Authors also thank Dr. Shilpy Sharma (Savitribai Phule Pune University, Pune) for active discussions during the data analysis and writing of the manuscript. HP is thankful to IISER Pune for the fellowship. HP is thankful to CSIR (India) [No. TG/10468/19-HRD] and Infosys Foundation [No. IISER-P/InfyFnd/Trv/139] for Travel support to present part of this work at Gordon Research Conference on Computational Aspects of Biomolecular Research, Switzerland, June 9-14, 2019.

## Supplementary Information

Supplementary Information (SI) available.

