## Supplementary Material for "Conformational dynamics at microsecond timescale in the RNA-binding regions of dsRNA-binding domains"

H Paithankar, and J Chugh

**Running Title**: Conformational dynamics in dsRNA-binding domains

**SUPPLEMENTARY FIGURES**

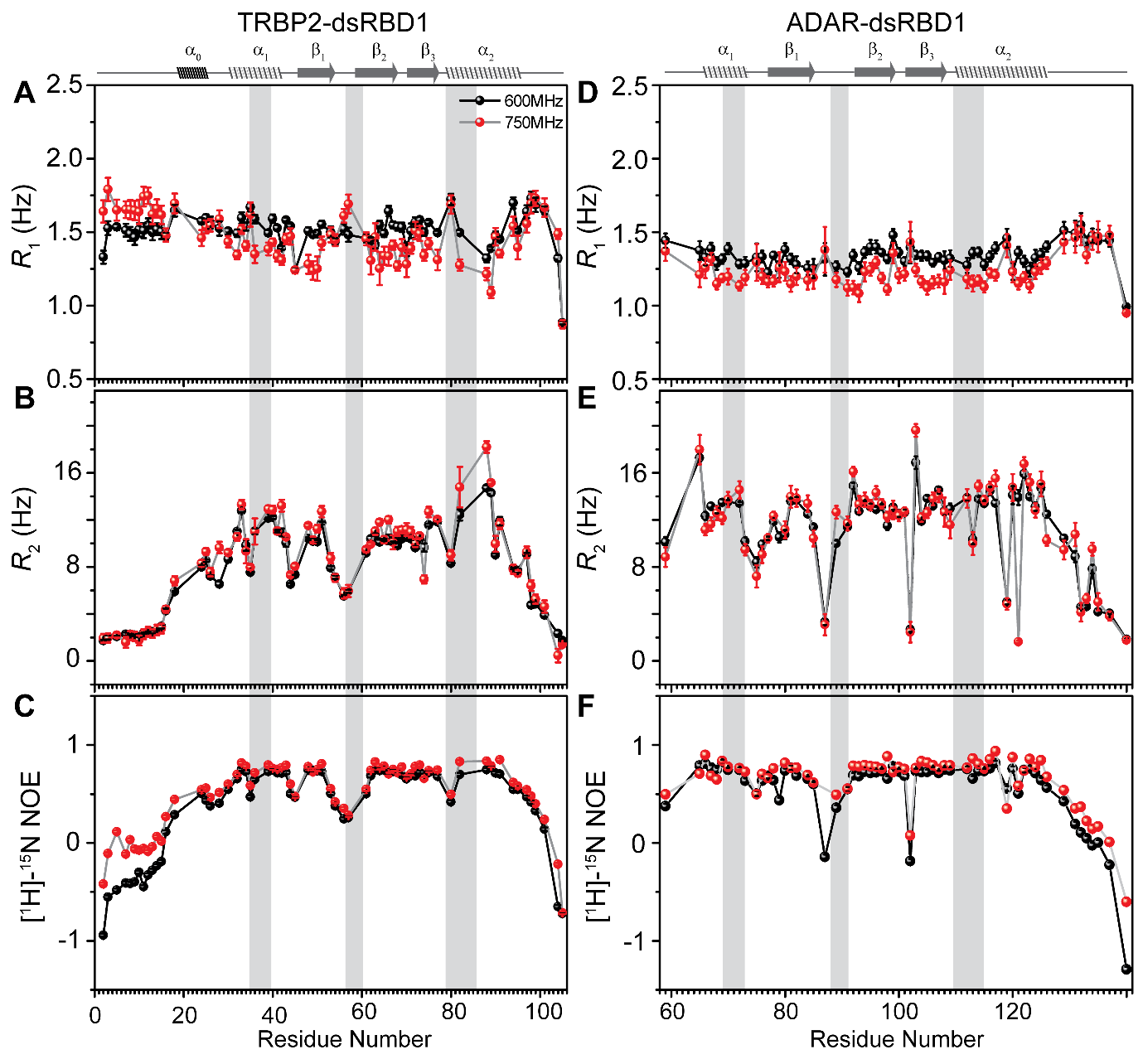

Figure S1: ^15^N-relaxation parameters ^15^N-R_1_ (A and D), ^15^N-R_2_ (B and E) and ^1^H-^15^N-nOe (C and F) for TRBP2-dsRBD1 and dADAR-dsRBD1 measured at 600 MHz (black) and at 750 MHz (red). The secondary structure of the two dsRBDs has been mentioned on the top, and the grey columns have been used to mark the RNA-binding region of the proteins.

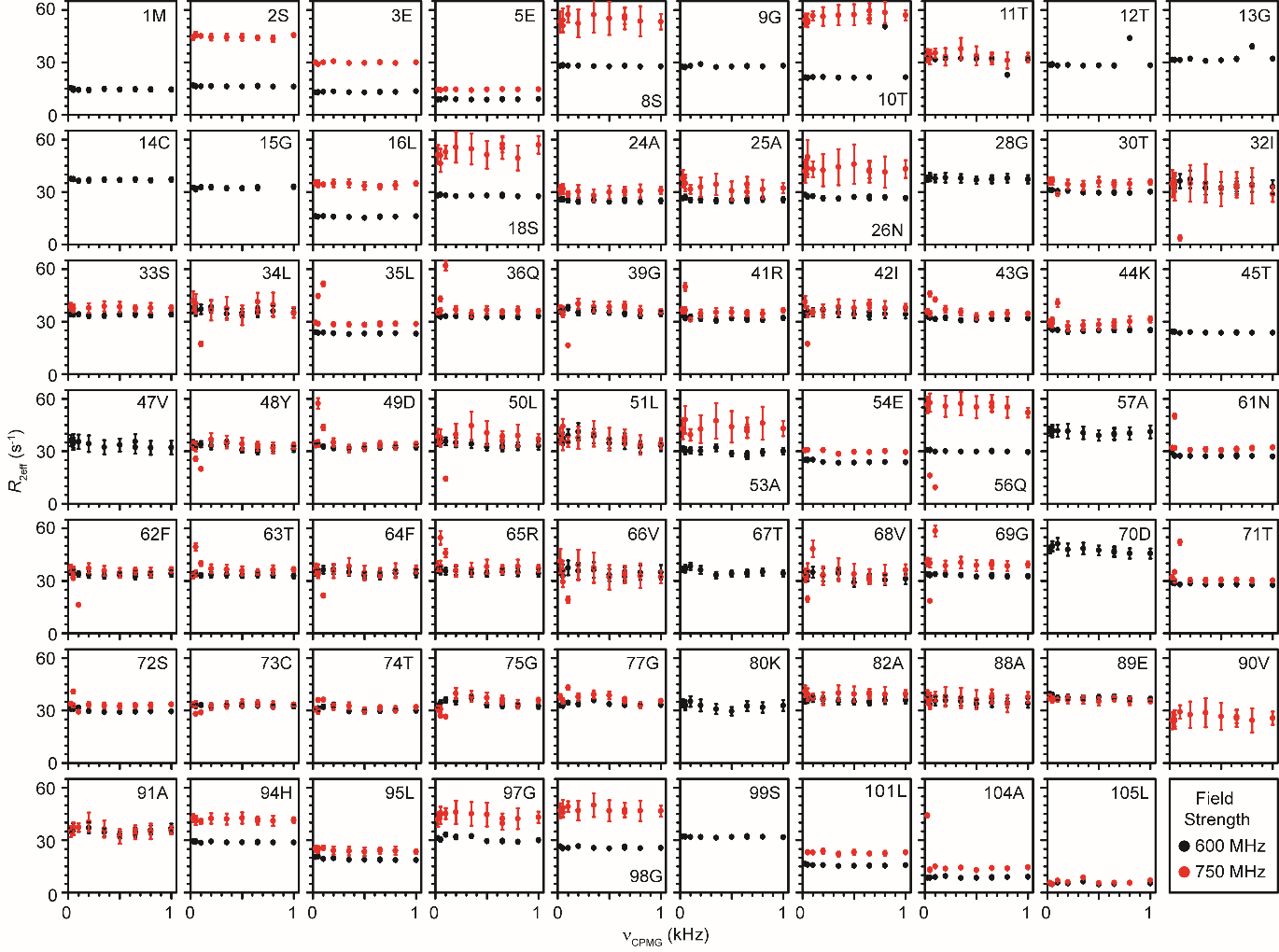

Figure S2: *R*_2eff_ rates as obtained from CPMG relaxation dispersion experiment plotted against the CPMG frequency for TRBP2-dsRBD1 measured at two magnetic fields 600 MHz (black) and 750 MHz (red).

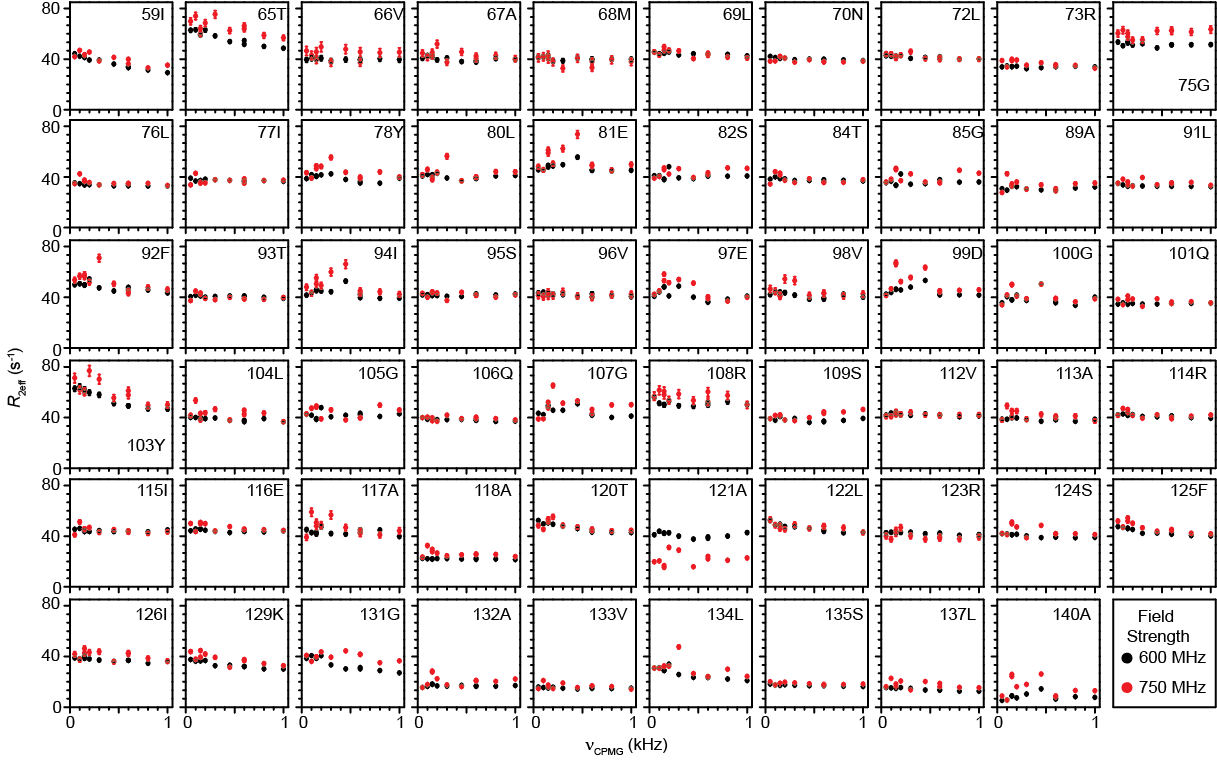

Figure S3: R_2eff_ rates as obtained from CPMG relaxation dispersion experiment plotted against the CPMG frequency for dADAR-dsRBD1 measured at two magnetic fields 600 MHz (black) and 750 MHz (red).

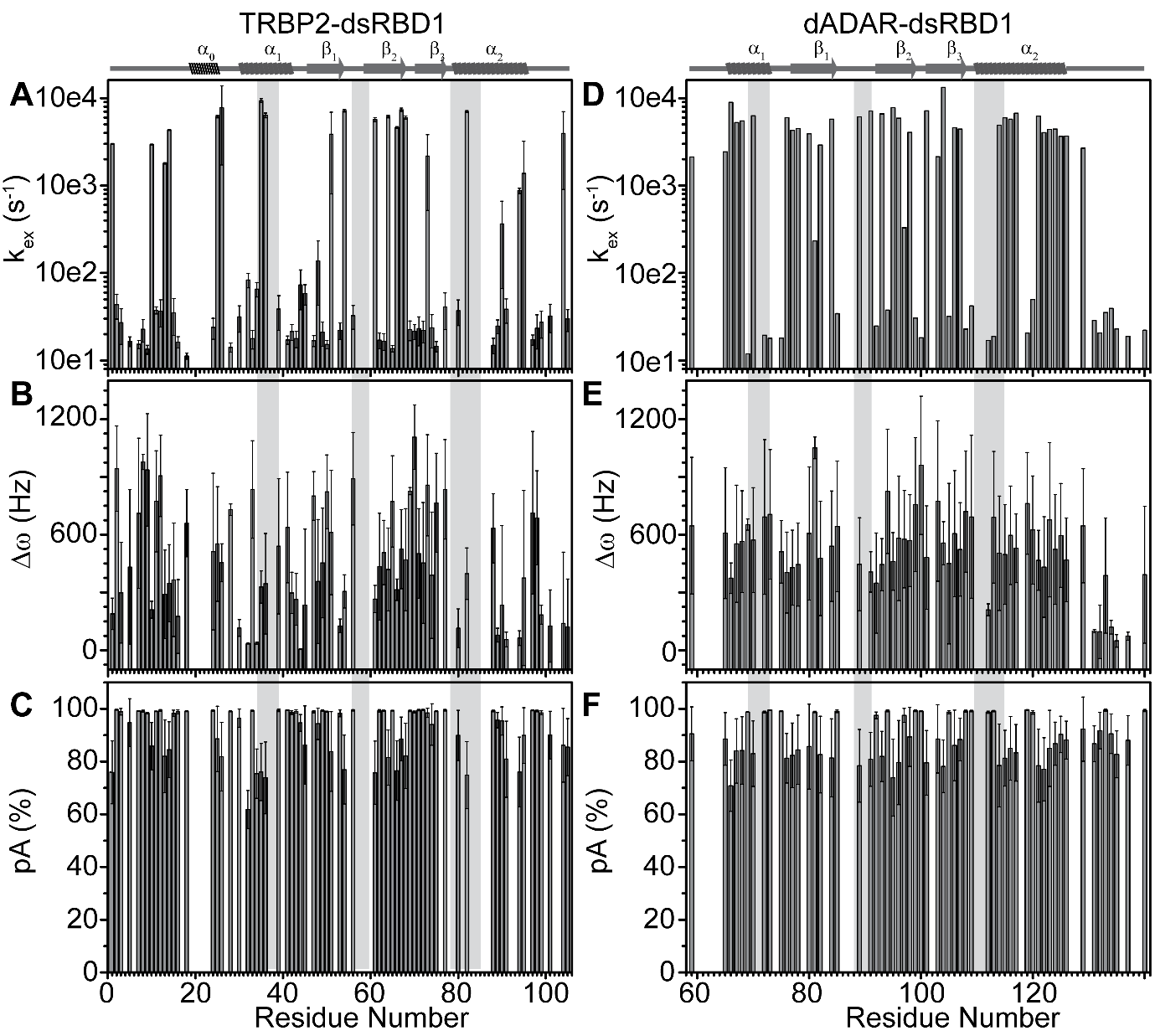

Figure S4: Dynamics parameters, rate of exchange between ground state and excited state (*k*_ex_), chemical shift difference between ground state and excited state (Δω), and population of the ground state (p_A_) as obtained by ‘Geometric approximation method’ from HARD experiments plotted against residue number for (A, B, C) TRBP2-dsRBD1 and (D, E, F) dADAR-dsRBD1, respectively. The secondary structure of the two dsRBDs has been mentioned on the top, and grey columns have been used to mark the RNA binding region of the proteins.

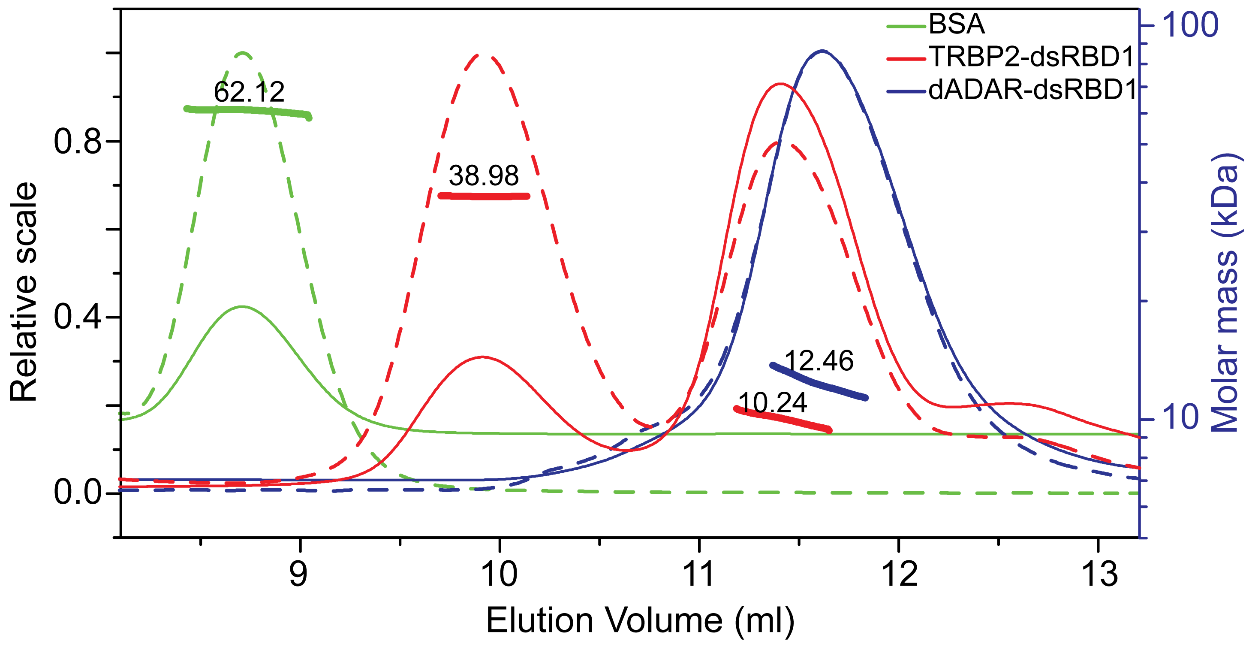

Figure S5: SEC-MALS analysis of TRBP2-dsRBD1 (red), dADAR-dsRBD1 (blue) proteins and BSA (Green) as reference protein. The axes on right indicates relative intensity for light scattering data (dashed line) and refractive index (solid line) for each protein. Molecular mass determined for the peak has been marked on the peak with scale on right of the plot. The reference protein BSA and dADAR-dsRBD1 showed a peak each with molecular mass of 62.12 kDa (±0.38%) and 12.46 kDa (±1.49%) respectively. The two peaks observed for TRBP2-dsRBD1 represent molecular mass of 38.98 kDa (±0.95%) and 10.24 kDa (±1.16%).

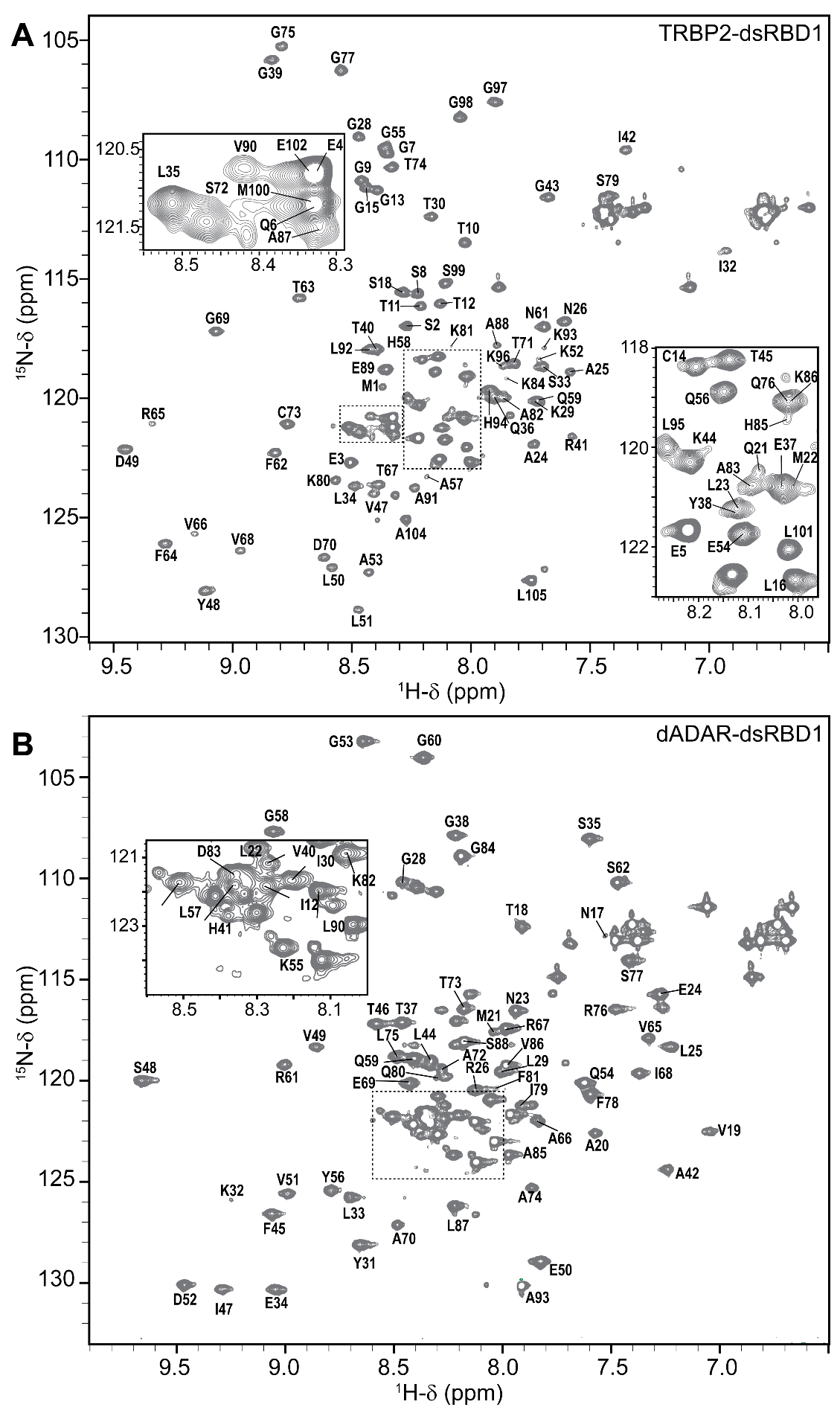

Figure S6: ^1^H-^15^N-HSQC spectrum for (A) TRBP2-dsRBD1 (reported previously by our group (1)) and (B) dADAR-dsRBD1 with resonance assignments (transferred from reported in literature (2) as described in methods section) at 25 °C in Buffer A, pH 6.4.

Table S1: Model-free parameters obtained for TRBP2-dsRBD1

| **Residue Number** | ***S^2^*** | | ***R_ex_* (Hz)** | | ***S^2^_f_*** | | ***S^2^_s_*** | | ***τ_e_* (ps)** | | ***τ_f_* (ps)** | | ***τ_s_* (ps)** | |
| --- | --- | --- | --- | --- | --- | --- | --- | --- | --- | --- | --- | --- | --- | --- |
|  | value | error | value | error | value | error | value | error | value | error | value | error | value | error |
| **2** | 0.01 | 0.01 |  |  | 0.73 | 0.04 | 0.02 | 0.01 |  |  | 100 | 23 | 757 | 47 |
| **3** | 0.01 | 0.01 |  |  | 0.77 | 0.04 | 0.01 | 0.02 |  |  | 96 | 23 | 921 | 53 |
| **5** | 0.00 | 0.01 | 0.25 | 0.12 | 0.83 | 0.01 |  |  |  |  |  |  | 891 | 13 |
| **7** | 0.03 | 0.01 |  |  | 0.58 | 0.03 | 0.05 | 0.02 |  |  | 118 | 10 | 1348 | 119 |
| **8** | 0.05 | 0.01 |  |  | 0.84 | 0.02 | 0.06 | 0.01 |  |  |  |  | 846 | 8 |
| **9** | 0.04 | 0.01 |  |  | 0.65 | 0.03 | 0.06 | 0.02 |  |  | 92 | 12 | 1145 | 87 |
| **10** | 0.01 | 0.01 |  |  | 0.51 | 0.02 | 0.03 | 0.01 |  |  | 105 | 7 | 1627 | 142 |
| **11** | 0.05 | 0.01 |  |  | 0.74 | 0.04 | 0.07 | 0.01 |  |  | 95 | 18 | 990 | 63 |
| **12** | 0.04 | 0.01 |  |  | 0.56 | 0.03 | 0.08 | 0.01 |  |  | 134 | 11 | 1533 | 141 |
| **13** | 0.02 | 0.01 |  |  | 0.54 | 0.02 | 0.10 | 0.01 |  |  | 101 | 9 | 1407 | 105 |
| **14** | 0.07 | 0.01 |  |  | 0.67 | 0.03 | 0.12 | 0.01 |  |  | 78 | 9 | 1200 | 83 |
| **15** | 0.06 | 0.01 |  |  | 0.56 | 0.03 | 0.18 | 0.02 |  |  | 98 | 8 | 1553 | 140 |
| **16** | 0.20 | 0.02 |  |  | 0.63 | 0.02 | 0.38 | 0.02 |  |  | 56 | 6 | 1529 | 153 |
| **18** | 0.24 | 0.08 | 1.12 | 0.84 | 0.73 | 0.06 | 0.36 | 0.10 |  |  | 43 | 11 | 1586 | 269 |
| **24** | 0.61 | 0.02 |  |  | 0.78 | 0.02 | 0.80 | 0.02 |  |  | 52 | 10 | 1506 | 770 |
| **25** | 0.56 | 0.03 | 0.03 | 0.35 | 0.90 | 0.02 | 0.66 | 0.03 |  |  |  |  | 1084 | 55 |
| **26** | 0.45 | 0.01 |  |  | 0.78 | 0.03 | 0.68 | 0.02 |  |  | 65 | 12 | 1623 | 279 |
| **28** | 0.00 | 0.05 | 4.33 | 0.78 | 0.54 | 0.04 |  |  |  |  | 25 | 5 | 2287 | 247 |
| **30** | 0.51 | 0.12 | 1.40 | 0.76 | 0.70 | 0.06 | 0.69 | 0.13 |  |  | 23 | 8 | 1685 | 1125 |
| **32** | 0.65 | 0.01 |  |  | 0.88 | 0.01 | 0.92 | 0.01 |  |  |  |  | 1574 | 76 |
| **33** | 0.73 | 0.01 |  |  |  |  |  |  | 2552 | 231 |  |  |  |  |
| **34** | 0.43 | 0.21 | 2.99 | 1.53 | 0.70 | 0.08 | 0.58 | 0.25 |  |  |  |  | 3121 | 1148 |
| **35** | 0.54 | 0.01 |  |  | 0.87 | 0.01 | 0.60 | 0.01 |  |  |  |  | 1156 | 40 |
| **36** | 0.65 | 0.10 |  |  | 0.91 | 0.06 | 0.84 | 0.08 |  |  |  |  | 1679 | 210 |
| **39** | 0.83 | 0.03 |  |  | 0.93 | 0.02 | 0.95 | 0.02 |  |  |  |  | 1457 | 265 |
| **40** | 0.60 | 0.03 |  |  | 0.91 | 0.02 | 0.90 | 0.02 |  |  |  |  | 2613 | 214 |
| **41** | 0.87 | 0.01 |  |  |  |  |  |  | 43 | 5 |  |  |  |  |
| **42** | 0.34 | 0.15 | 5.11 | 1.11 | 0.62 | 0.06 | 0.51 | 0.20 |  |  |  |  | 2933 | 743 |
| **43** | 0.35 | 0.04 |  |  | 0.77 | 0.02 | 0.75 | 0.04 |  |  |  |  | 3037 | 234 |
| **44** | 0.23 | 0.01 |  |  | 0.68 | 0.02 | 0.61 | 0.01 |  |  | 21 | 6 | 2179 | 268 |
| **45** | 0.25 | 0.02 |  |  | 0.59 | 0.02 | 0.71 | 0.02 |  |  | 31 | 3 | 2434 | 299 |
| **48** | 0.79 | 0.01 | 0.61 | 0.14 |  |  |  |  | 16 | 3 |  |  |  |  |
| **49** | 0.76 | 0.01 |  |  |  |  |  |  | 20 | 3 |  |  |  |  |
| **50** | 0.77 | 0.01 | 0.21 | 0.16 |  |  |  |  | 19 | 3 |  |  |  |  |
| **51** | 0.61 | 0.05 |  |  | 0.88 | 0.03 | 0.87 | 0.04 |  |  |  |  | 2836 | 332 |
| **53** | 0.42 | 0.01 |  |  | 0.74 | 0.02 | 0.73 | 0.02 |  |  | 42 | 7 | 2264 | 444 |
| **54** | 0.48 | 0.01 |  |  | 0.69 | 0.02 | 0.68 | 0.02 |  |  | 54 | 7 | 1623 | 405 |
| **56** | 0.32 | 0.01 |  |  | 0.69 | 0.03 | 0.47 | 0.02 |  |  | 63 | 8 | 1630 | 247 |
| **57** | 0.43 | 0.02 |  |  | 0.65 | 0.02 | 0.56 | 0.03 |  |  | 87 | 8 | 2077 | 812 |
| **61** | 0.64 | 0.01 |  |  | 0.80 | 0.02 | 0.83 | 0.02 |  |  | 50 | 12 | 1369 | 454 |
| **62** | 0.43 | 0.01 |  |  | 0.75 | 0.01 | 0.89 | 0.01 |  |  |  |  | 2367 | 138 |
| **63** | 0.80 | 0.01 |  |  |  |  |  |  | 8 | 3 |  |  |  |  |
| **64** | 0.72 | 0.01 |  |  | 0.85 | 0.01 | 0.94 | 0.02 |  |  |  |  | 1825 | 220 |
| **65** | 0.77 | 0.01 |  |  |  |  |  |  | 14 | 3 |  |  |  |  |
| **66** | 0.83 | 0.01 | 0.03 | 0.15 |  |  |  |  | 42 | 6 |  |  |  |  |
| **67** | 0.68 | 0.09 | 0.71 | 0.88 | 0.82 | 0.04 | 0.80 | 0.07 |  |  |  |  | 1719 | 501 |
| **68** | 0.80 | 0.01 | 0.04 | 0.14 |  |  |  |  | 31 | 3 |  |  |  |  |
| **69** | 0.77 | 0.04 |  |  | 0.85 | 0.02 | 0.92 | 0.03 |  |  |  |  | 1072 | 293 |
| **70** | 0.81 | 0.01 |  |  |  |  |  |  | 45 | 4 |  |  |  |  |
| **71** | 0.86 | 0.01 |  |  |  |  |  |  | 56 | 7 |  |  |  |  |
| **72** | 0.70 | 0.04 |  |  | 0.84 | 0.02 | 0.88 | 0.04 |  |  |  |  | 1608 | 258 |
| **73** | 0.81 | 0.03 |  |  | 0.86 | 0.02 | 0.96 | 0.03 |  |  |  |  | 993 | 367 |
| **74** | 0.49 | 0.04 |  |  | 0.69 | 0.01 | 0.83 | 0.05 |  |  | 25 | 4 | 6605 | 2360 |
| **75** | 0.73 | 0.07 | 1.54 | 0.74 | 0.84 | 0.03 | 0.84 | 0.05 |  |  |  |  | 1283 | 464 |
| **77** | 0.86 | 0.01 |  |  |  |  |  |  | 46 | 5 |  |  |  |  |
| **80** | 0.52 | 0.01 |  |  | 0.84 | 0.03 | 0.71 | 0.02 |  |  | 98 | 18 | 1706 | 365 |
| **82** | 0.32 | 0.25 | 6.61 | 2.03 | 0.63 | 0.09 | 0.45 | 0.33 |  |  |  |  | 3985 | 1712 |
| **88** | 0.61 | 0.13 | 5.17 | 1.29 | 0.75 | 0.06 | 0.84 | 0.12 |  |  |  |  | 2925 | 1309 |
| **89** | 0.75 | 0.01 | 3.45 | 0.13 |  |  |  |  | 10 | 2 |  |  |  |  |
| **90** | 0.74 | 0.01 |  |  |  |  |  |  |  |  |  |  |  |  |
| **91** | 0.56 | 0.14 | 2.28 | 1.12 | 0.74 | 0.06 | 0.69 | 0.14 |  |  |  |  | 2661 | 932 |
| **94** | 0.53 | 0.01 |  |  | 0.79 | 0.02 | 0.67 | 0.02 |  |  | 36 | 12 | 1966 | 640 |
| **95** | 0.42 | 0.02 |  |  | 0.70 | 0.02 | 0.74 | 0.03 |  |  | 36 | 5 | 2920 | 1135 |
| **97** | 0.66 | 0.01 |  |  | 0.85 | 0.03 | 0.74 | 0.03 |  |  | 55 | 23 | 1049 | 375 |
| **98** | 0.00 | 0.08 | 1.95 | 0.67 | 0.58 | 0.04 |  |  |  |  | 44 | 6 | 3521 | 594 |
| **99** | 0.10 | 0.01 |  |  | 0.63 | 0.02 | 0.37 | 0.01 |  |  | 66 | 6 | 2383 | 274 |
| **101** | 0.13 | 0.01 |  |  | 0.59 | 0.02 | 0.28 | 0.02 |  |  | 84 | 5 | 2246 | 282 |
| **104** | 0.02 | 0.01 |  |  | 0.64 | 0.03 | 0.03 | 0.02 |  |  | 74 | 9 | 865 | 43 |
| **105** | 0.01 | 0.01 |  |  | 0.21 | 0.00 | 0.07 | 0.04 |  |  | 70 | 1 | 3760 | 473 |

Table S2: Model-free parameters for dADAR-dsRBD1

| **Residue Number** | ***S^2^*** | | ***R_ex_* (Hz)** | | ***S^2^_f_*** | | ***S^2^_s_*** | | ***τ_e_* (ps)** | | ***τ_f_* (ps)** | | ***τ_s_* (ps)** | |
| --- | --- | --- | --- | --- | --- | --- | --- | --- | --- | --- | --- | --- | --- | --- |
|  | value | error | value | error | value | error | value | error | value | error | value | error | value | error |
| **59** | 0.62 | 0.03 |  |  | 0.92 | 0.02 | 0.70 | 0.02 |  |  |  |  | 874 | 61 |
| **65** | 0.95 | 0.03 | 3.56 | 0.43 |  |  |  |  | 91 | 326 |  |  |  |  |
| **66** | 0.77 | 0.03 |  |  | 0.86 | 0.02 | 0.93 | 0.04 |  |  |  |  | 4235 | 2292 |
| **67** | 0.85 | 0.02 |  |  | 0.95 | 0.02 | 0.92 | 0.02 |  |  |  |  | 1398 | 357 |
| **68** | 0.89 | 0.01 |  |  |  |  |  |  | 64 | 16 |  |  |  |  |
| **69** | 0.94 | 0.01 |  |  |  |  |  |  |  |  |  |  |  |  |
| **70** | 0.95 | 0.02 |  |  |  |  |  |  | 94 | 102 |  |  |  |  |
| **72** | 0.90 | 0.02 | 0.74 | 0.27 |  |  |  |  | 43 | 12 |  |  |  |  |
| **73** | 0.63 | 0.02 |  |  | 0.81 | 0.02 | 0.81 | 0.02 |  |  |  |  | 1354 | 138 |
| **75** | 0.49 | 0.06 |  |  | 0.69 | 0.03 | 0.74 | 0.07 |  |  | 48 | 13 | 3040 | 2513 |
| **76** | 0.63 | 0.03 |  |  | 0.81 | 0.02 | 0.80 | 0.02 |  |  |  |  | 1381 | 144 |
| **77** | 0.70 | 0.01 |  |  | 0.82 | 0.01 | 0.88 | 0.01 |  |  |  |  | 1127 | 144 |
| **78** | 0.81 | 0.01 |  |  | 0.91 | 0.01 | 0.92 | 0.01 |  |  |  |  | 841 | 118 |
| **80** | 0.71 | 0.03 |  |  | 0.84 | 0.02 | 0.87 | 0.02 |  |  |  |  | 2022 | 589 |
| **81** | 0.91 | 0.02 | 0.65 | 0.35 |  |  |  |  | 39 | 18 |  |  |  |  |
| **82** | 0.85 | 0.11 | 1.19 | 1.25 | 0.92 | 0.07 | 0.92 | 0.06 |  |  |  |  | 799 | 584 |
| **84** | 0.86 | 0.02 | 0.15 | 0.29 |  |  |  |  | 66 | 15 |  |  |  |  |
| **85** | 0.80 | 0.01 |  |  |  |  |  |  | 56 | 8 |  |  |  |  |
| **89** | 0.41 | 0.09 | 3.41 | 1.03 | 0.73 | 0.05 | 0.57 | 0.09 |  |  |  |  | 1063 | 145 |
| **91** | 0.80 | 0.01 |  |  |  |  |  |  | 72 | 8 |  |  |  |  |
| **92** | 0.92 | 0.02 | 1.42 | 0.31 |  |  |  |  | 80 | 57 |  |  |  |  |
| **93** | 0.79 | 0.10 | 1.20 | 1.18 | 0.86 | 0.07 | 0.92 | 0.06 |  |  |  |  | 803 | 624 |
| **94** | 0.94 | 0.01 | 0.02 | 0.19 |  |  |  |  | 80 | 43 |  |  |  |  |
| **95** | 0.88 | 0.02 |  |  | 0.97 | 0.01 | 0.93 | 0.01 |  |  |  |  | 1089 | 225 |
| **96** | 0.75 | 0.11 | 1.65 | 1.16 | 0.90 | 0.06 | 0.84 | 0.06 |  |  |  |  | 1709 | 504 |
| **97** | 0.92 | 0.01 |  |  |  |  |  |  | 67 | 16 |  |  |  |  |
| **98** | 0.77 | 0.02 |  |  | 0.87 | 0.01 | 0.91 | 0.01 |  |  |  |  | 1126 | 197 |
| **99** | 0.94 | 0.01 |  |  |  |  |  |  | 84 | 98 |  |  |  |  |
| **100** | 0.83 | 0.02 |  |  | 0.91 | 0.01 | 0.94 | 0.01 |  |  |  |  | 1107 | 241 |
| **101** | 0.87 | 0.01 |  |  |  |  |  |  | 55 | 10 |  |  |  |  |
| **102** | 0.08 | 0.02 |  |  | 0.59 | 0.06 | 0.13 | 0.03 |  |  | 62 | 16 | 1247 | 196 |
| **103** | 0.66 | 0.16 | 5.91 | 1.47 | 0.81 | 0.09 | 0.81 | 0.13 |  |  |  |  | 1917 | 825 |
| **104** | 0.82 | 0.01 |  |  | 0.90 | 0.01 | 0.94 | 0.01 |  |  |  |  | 1231 | 275 |
| **105** | 0.92 | 0.01 |  |  |  |  |  |  | 56 | 17 |  |  |  |  |
| **106** | 0.89 | 0.01 | 0.32 | 0.25 |  |  |  |  | 35 | 11 |  |  |  |  |
| **107** | 0.91 | 0.02 | 0.88 | 0.32 |  |  |  |  | 70 | 31 |  |  |  |  |
| **108** | 0.90 | 0.02 |  |  |  |  |  |  | 38 | 34 |  |  |  |  |
| **109** | 0.91 | 0.02 |  |  |  |  |  |  | 49 | 77 |  |  |  |  |
| **112** | 0.91 | 0.03 | 0.75 | 0.42 |  |  |  |  | 46 | 68 |  |  |  |  |
| **113** | 0.68 | 0.03 |  |  | 0.83 | 0.02 | 0.84 | 0.02 |  |  |  |  | 1546 | 229 |
| **114** | 0.86 | 0.08 | 1.14 | 0.87 | 0.93 | 0.05 | 0.94 | 0.04 |  |  |  |  | 1075 | 665 |
| **115** | 0.90 | 0.02 | 0.17 | 0.23 |  |  |  |  | 51 | 14 |  |  |  |  |
| **116** | 0.85 | 0.11 | 1.41 | 1.14 | 0.92 | 0.06 | 0.92 | 0.07 |  |  |  |  | 1641 | 1408 |
| **117** | 0.97 | 0.01 |  |  |  |  |  |  |  |  |  |  |  |  |
| **120** | 0.64 | 0.28 | 3.46 | 2.62 | 0.77 | 0.15 | 0.83 | 0.29 |  |  |  |  | 2614 | 2205 |
| **122** | 0.94 | 0.02 | 1.99 | 0.35 |  |  |  |  | 87 | 85 |  |  |  |  |
| **123** | 0.82 | 0.12 | 2.12 | 1.28 | 0.87 | 0.08 | 0.94 | 0.07 |  |  |  |  | 1459 | 1547 |
| **124** | 0.88 | 0.02 |  |  | 0.95 | 0.02 | 0.94 | 0.02 |  |  |  |  | 957 | 266 |
| **125** | 0.62 | 0.17 | 4.82 | 1.91 | 0.81 | 0.10 | 0.76 | 0.15 |  |  |  |  | 1648 | 577 |
| **126** | 0.76 | 0.02 |  |  | 0.94 | 0.01 | 0.84 | 0.02 |  |  |  |  | 926 | 88 |
| **129** | 0.69 | 0.02 |  |  |  |  |  |  | 922 | 59 |  |  |  |  |
| **131** | 0.60 | 0.03 |  |  |  |  |  |  | 763 | 53 |  |  |  |  |
| **132** | 0.22 | 0.01 |  |  | 0.81 | 0.03 | 0.28 | 0.02 |  |  |  |  | 1059 | 27 |
| **133** | 0.24 | 0.01 |  |  | 0.65 | 0.05 | 0.38 | 0.02 |  |  | 60 | 16 | 1354 | 275 |
| **134** | 0.29 | 0.12 | 2.37 | 1.26 | 0.69 | 0.10 | 0.42 | 0.14 |  |  | 91 | 40 | 1257 | 468 |
| **135** | 0.19 | 0.02 |  |  | 0.62 | 0.05 | 0.32 | 0.03 |  |  | 72 | 15 | 1475 | 333 |
| **137** | 0.18 | 0.01 |  |  | 0.66 | 0.06 | 0.29 | 0.03 |  |  | 93 | 19 | 1153 | 213 |
| **140** | 0.05 | 0.00 |  |  | 0.68 | 0.00 | 0.08 | 0.00 |  |  |  |  | 505 | 6 |
